# Phylogenetic insights into the origin of piroplasmid infection in endemic carnivores of Madagascar

**DOI:** 10.64898/2026.11.12.766023

**Authors:** Fidisoa T. Rasambainarivo, Kayla J. Evans, Olivier A. Rasolofoniaina, Santatriniaina F. Randrianarisoa, Ananías A. Escalante, M. Andreína Pacheco, Zea Walton, Benjamin L. Rice

**Author notes:** Corresponding author: Fidisoa Rasambainarivo, Department of Biology, East Carolina University, Greenville, NC, USA.

## Abstract

Piroplasmids (order Piroplasmida), including the genera *Babesia*, *Theileria*, and *Cytauxzoon*, are tick-borne protozoan parasites of medical, veterinary, and economic importance. Understanding the diversity and evolutionary history of piroplasmids infecting endangered wildlife can provide insights into their origins, natural hosts, and potential to cause severe disease. Using nuclear *18S rRNA* and mitochondrial *cytb* gene sequences, we characterized piroplasmid parasites infecting wild carnivorans across two protected areas in Madagascar. We detected piroplasmid infections in 91.5% of sampled individuals (n = 106; 95% CI: 84.5–96%) across three endemic carnivoran species, with infections associated with distinct, potentially host-associated *Babesia* sp. and *Cytauxzoon* sp. lineages. Parasites detected in broad-striped vontsira (*Galidictis fasciata*) were most closely related to *Babesia* sp. previously reported from lemurs and fruit bats in Madagascar, whereas parasites detected in fosa (*Cryptoprocta ferox*) and ring-tailed vontsira (*Galidia elegans*) were more closely related to *Cytauxzoon* lineages reported from carnivores elsewhere, including meerkats in South Africa. Phylogenetic analyses revealed partial incongruence between the two genetic markers: *18S rRNA* sequences placed the parasites into two distinct lineages, whereas the mitochondrial *cytb* phylogeny grouped them into a single, well-supported clade. This may reflect shared ancestry, although methodological artifacts and differences in taxon sampling cannot be excluded. Our findings indicate that endemic Malagasy carnivorans harbor distinct piroplasmid lineages, potentially including locally evolved and previously undescribed parasites.

## Introduction

Piroplasmids (order Piroplasmida) are unicellular tick-borne protozoan parasites comprising species from the genera *Babesia*, *Theileria,* and *Cytauxzoon*. Piroplasmid infections (piroplasmosis) occur in a wide range of domestic and wild animals and in humans, causing diseases of medical, economic, and veterinary importance worldwide (Schnittger et al. 2012, 2022; Solano-Gallego et al. 2016). Piroplasmids are among the most prevalent blood parasites of mammals, and several can cross species barriers to infect alternative vertebrate hosts, often with severe clinical consequences (Leemans et al. 1999; Zintl et al. 2003; Schnittger et al. 2012). For example, human infection by *Babesia divergens*, whose natural reservoirs are bovids, requires rapid diagnosis as infection frequently results in major organ failure and death within a few days (Kjemtrup and Conrad 2000). Concerns about zoonotic *Babesia* sp. infections have grown (Yabsley 2007). In general, the severity of illness depends on multiple factors, including the immunocompetence of the infected host and whether the triad of ticks, host, and parasite has co-evolved (Florin-Christensen and Schnittger 2009; Penzhorn 2011).

Piroplasmids are also increasingly recognized as important wildlife pathogens (Shock et al. 2012, 2024; Yabsley and Shock 2012). In wild mammals, infections with *Babesia*, *Theileria*, and *Cytauxzoon* can cause substantial morbidity and mortality, particularly after spillover into immunologically naïve hosts, or in the presence of coinfection and may further stress vulnerable populations (Penzhorn 2006; Munson et al. 2008). For example, unusually high mortality following canine distemper virus (CDV) outbreaks in African lions in 1994 and 2001 was attributed to co-infections with *Babesia* sp., whereas CDV alone was not associated with measurable mortality in lion populations (Munson et al. 2008). Consequently, piroplasmids represent an emerging concern in wildlife health and conservation, particularly in regions experiencing rapid environmental change and intensifying anthropogenic pressures (Munson et al. 2008). Yet, the diversity, host associations, and evolutionary origins of piroplasmids infecting wildlife remain poorly characterized (Soares et al. 2017). Distinguishing long-standing, potentially coevolved host-parasite associations from recent spillovers is critical for understanding disease risk and informing wildlife conservation.

Phylogenetic inference of piroplasmid relationships commonly combines the nuclear 18S ribosomal RNA (*18S rRNA*) gene with the mitochondrial cytochrome b (*cytb*) gene (Kjemtrup et al. 2000; Lack et al. 2012). The *18S rRNA* gene is highly conserved and evolves relatively slowly, making it useful for establishing a broad phylogenetic framework and resolving relationships among divergent piroplasmid lineages. However, its high degree of conservation can limit the ability to distinguish closely related or recently diverged taxa. On the other hand, mitochondrial markers such as *cytb* gene have been informative for resolving more recent divergences in Apicomplexans (the phylum containing piroplasmids) (Escalante et al. 1998; Pacheco and Escalante 2023). However, mitochondrial markers are correspondingly prone to substitution saturation at deeper divergence times, where accumulated multiple substitutions can erode phylogenetic signals (Nguyen and Ho 2016; Pacheco et al. 2018a; Cuadrado et al. 2024). Thus, these markers provide complementary information, with 18S rRNA contributing to the broader evolutionary framework and the *cytb* gene providing insight into relationships among closely related piroplasmids.

Madagascar provides a particularly informative system for investigating the evolutionary origins and host associations of piroplasmids. The Eupleridae comprise a unique family of carnivorans consisting of seven monospecific genera, all endemic to Madagascar, that have evolved in geographic isolation from other carnivoran lineages for more than 20 million years (Yoder et al. 2003; Veron et al. 2017; Veron 2020). In addition to these endemic carnivorans, Madagascar now has several introduced carnivore species, including domestic dogs, domestic cats, and the Indian civet (*Viverricula indica*). These introduced carnivorans colonized the island relatively recently, approximately 2,000 years ago, and camera-trap studies suggest that they may increasingly overlap with and potentially replace native carnivorans, particularly in forests surrounding human settlements (Ardalan et al. 2015; Farris et al. 2017; Rasambainarivo et al. 2017; Sauther et al. 2020; Hixon et al. 2021). Such sympatry creates opportunities for pathogen transmission between introduced and endemic hosts. Indeed, frequent interactions among humans, domestic carnivores, and wild carnivores in Madagascar have been identified as potential pathways for pathogen transmission among sympatric species (Rasambainarivo et al. 2018).

Despite the potential importance of these interactions, the diversity and evolutionary origins of hemoparasites circulating among Madagascar’s endemic carnivorans remain poorly understood. Although researchers have documented piroplasmid infections in numerous wild carnivoran species worldwide, they have not previously described infections in Eupleridae or their sympatric domestic carnivorans in Madagascar (Alvarado-Rybak et al. 2016). Nevertheless, researchers have detected piroplasmids in other Malagasy wildlife, including lemurs and fruit bats, which harbor genetically distinct *Babesia* lineages (Larsen et al. 2016; Ranaivoson et al. 2019). Furthermore, critically endangered Verreaux’s sifaka (*Propithecus verreauxi)* in southwestern Madagascar were reported to harbor a piroplasmid lineage indistinguishable from *Babesia canis* detected in domestic dogs, suggesting that introduced piroplasmids may infect native Malagasy wildlife (Springer et al. 2015). These observations raise two alternative possibilities for piroplasmids infecting endemic Malagasy carnivorans: they may represent endemic lineages that diversified in Madagascar (prior to human settlement), or they may have been introduced through contemporary spillover transmission from domestic or other introduced carnivores.

Our first objective was therefore to investigate the evolutionary origins of piroplasmids infecting carnivorans in Madagascar. We hypothesized that piroplasmids infecting endemic Malagasy carnivorans reflect two distinct evolutionary origins: (i) locally evolved, endemic lineages shaped by Madagascar’s long-term geographic isolation, and (ii) recently introduced lineages resulting from contemporary spillover from domestic carnivorans at wildlife - domestic animal interfaces. Under hypothesis 1, local evolution and ancient diversification on Madagascar, we predicted that piroplasmid sequences from Eupleridae would form well-supported monophyletic or geographically structured clades distinct from known continental lineages and show closest affinities to other piroplasmids infecting endemic Malagasy wildlife. Under hypothesis 2, recent introduction and spillover, we predicted that sequences from endemic carnivorans would cluster closely with, or be nested within, lineages previously detected in domestic dogs, cats, or other introduced carnivore hosts, consistent with recent host switching at wildlife–domestic animal interfaces. We evaluated these alternatives using phylogenetic analyses of the *18S rRNA* and mitochondrial *cytb* genes. Our second objective was to assess potential cross-species transmission of piroplasmids among sympatric Eupleridae using the mitochondrial *cytb* marker.

## Methods

### Capture and specimen collection

We investigated piroplasmid infection in domestic and wild carnivorans living in or near the Betampona Natural Reserve (BET) in eastern Madagascar and Kirindy Mitea National Park (KMN) in southwestern Madagascar. BET encompasses 2,228 ha of protected lowland rainforest and is isolated from surrounding forest patches by agricultural land and human settlements. KMNP encompasses approximately 152,000 ha of dry deciduous forest in western Madagascar and contains some of the largest remaining contiguous tracts of dry forest in the country (Fig. 1).

**Figure 1.**
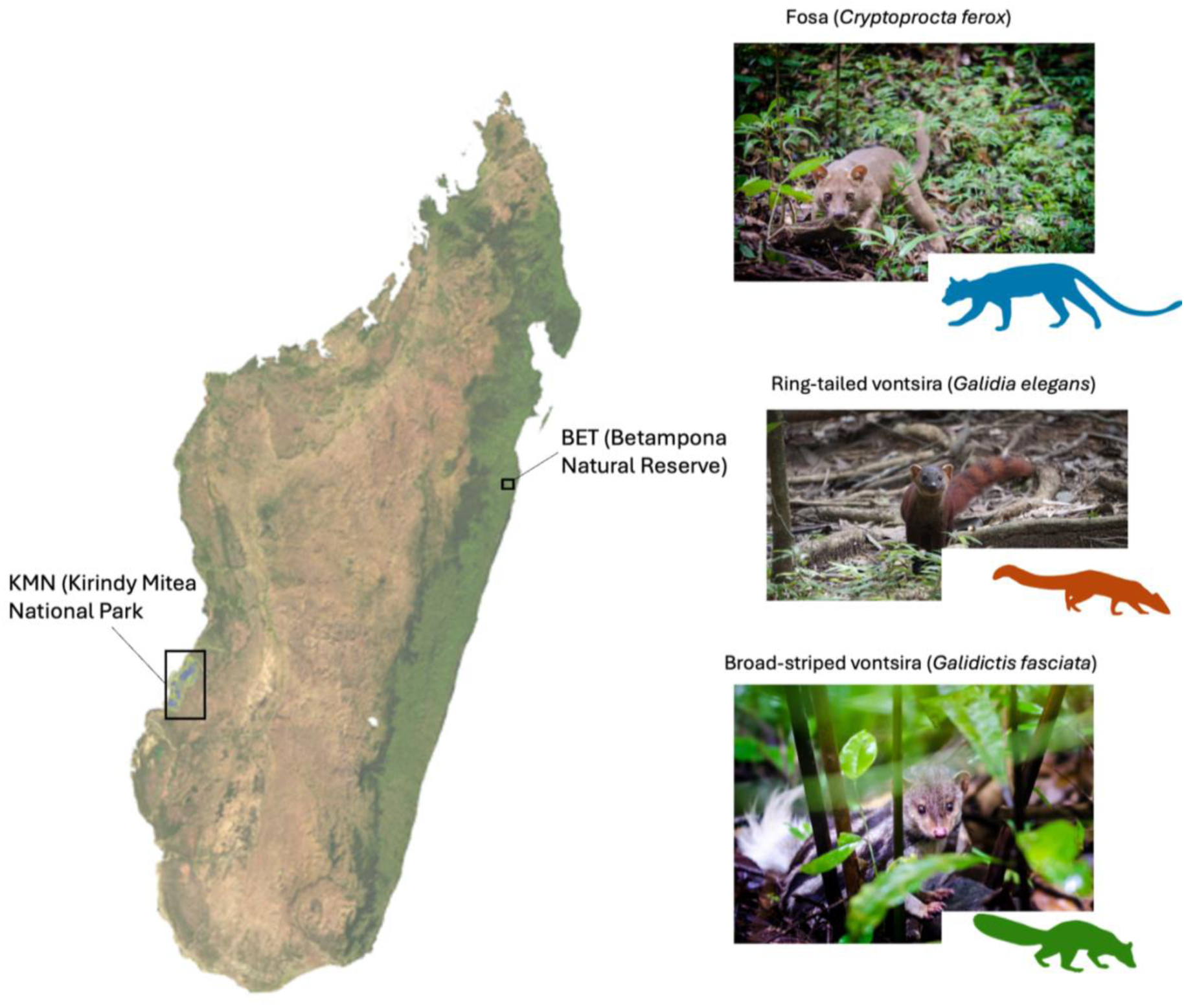
Locations of field sites and species surveyed in Madagascar.

All animal procedures were approved by the East Carolina University and Princeton University Institutional Animal Care and Use Committee (# 3064 [Princeton University], AUP #D380 [East Carolina University] and AUP # 00143 [University of Texas Austin]) and conducted in accordance with applicable research and animal welfare requirements in Madagascar under research permits # N125/22/MEDD/SG/DGGE/DAPRNE/SCBE, 175/23/MEDD/SG/DGGE/DAPRNE/SCBE, N155/24/MEDD/SG/DGGE/DAPRNE/SCBE (Rasambainarivo), and N°131/22/MEDD/SG/DGGE/DAPRNE/SCBE, N°131/23/MEDD/SG/DGGE/DAPRNE/SCBE, N°350/23/MEDD/SG/DGGE/DAPRNE/SCBE (Walton).

From June to July each year between 2022 and 2024, wild carnivorans were captured using Tomahawk live traps (Model 209, Tomahawk Live Trap, Tomahawk, Wisconsin, United States) baited with water, cat food, or chickens. Traps were checked twice a day (dawn and dusk). Captured animals were anesthetized by intramuscular injection of ketamine (approximately 4 mg/kg) and dexmedetomidine (0.01 mg/kg). We sampled domestic dogs and cats from villages surrounding BET with owner consent. For each animal, species, sex, and age category were recorded. Animals were classified as immature or adult based on body size, dentition, and reproductive condition (Rasambainarivo et al. 2025).

Approximately 2 mL of whole blood was collected by venipuncture of the femoral, cephalic or lateral saphenous vein into EDTA tubes and frozen until further processing. Following complete recovery from anesthesia, typically within 4 h, wild animals were released at their capture location. Ectoparasites were manually removed and preserved in 95% ethanol for subsequent identification. Ectoparasites were morphologically identified by Dr. Hans Klompen at Ohio State University and Dr Dmitry Apanaskevich at Georgia Southern University using published taxonomic keys (Hoogstraal, 1953).

### Laboratory analysis

Total DNA was extracted from 200 µL of whole blood using the DNeasy Blood & Tissue Kit (Qiagen) according to the manufacturer’s instructions. Portions of the nearly complete 18S rRNA (∼1,700 bp) and cytochrome b (*cytb*, ∼1,100 bp) genes were amplified to detect and genetically characterize piroplasmids infecting carnivorans in Madagascar. Each 50 μL reaction contained 1 μL of DNA template, 50 pmol of each primer, 10 nmol dNTPs, 75 nmol of MgCl2, 3.75 U AmpliTaq Gold DNA polymerase and a 1X concentration of GeneAmp PCR Gold Buffer (Applied Biosystems, Carlsbad, CA). Thermal cycling conditions consisted of an initial denaturation at 94°C for 5 minutes, followed by 35 – 40 amplification cycles (Table 1) and a final extension step at 72 C for 7 minutes. Primer sequences and PCR conditions are provided in Table 1. Amplicons were detected in 1% agarose gel stained with GelGreen Nucleic Acid Stain (Amplyus, Cambridge, MA, US). Results are reported with absolute and relative frequencies and 95% confidence intervals were calculated with the Wilson score method (Wilson 1927). Amplicons of the expected size were purified and submitted for Sanger sequencing at GENEWIZ from Azenta Life Sciences (South Plainfield, NJ, USA).

**Table 1:**
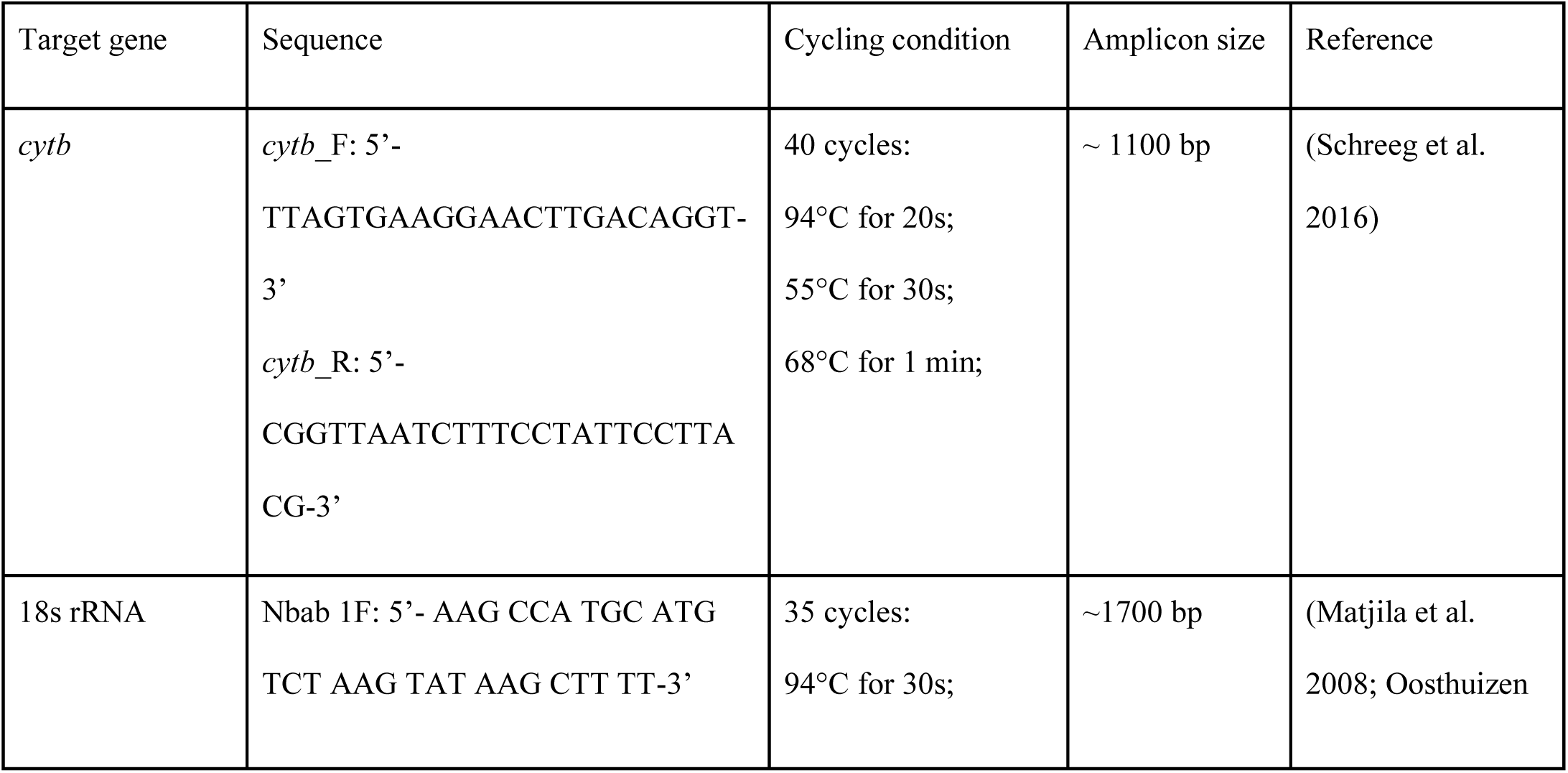
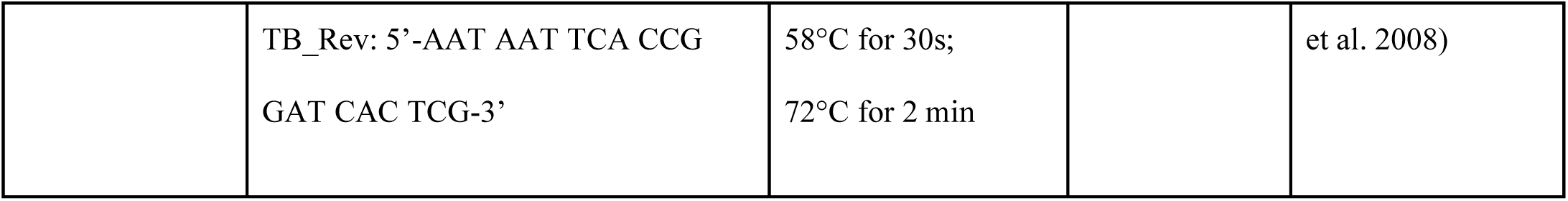
PCR primers and cycling conditions used to amplify the *cytb* and 18S rRNA genes in carnivores of Madagascar.

### Phylogenetic analyses

Sequences were assembled and edited using Geneious Prime version v. 2026.1.2 (Biomatters Ltd., New Zealand). Orthologous reference sequences were retrieved from GenBank and combined with sequences generated in this study (Table S1). Multiple sequence alignments were performed using MUSCLE in Geneious Prime with default settings and subsequently trimmed to uniform lengths of 1,479 bp (1,408 without gaps) for 18S rRNA and 1,170 bp (1,066 without gaps) for *cytb* gene. Bayesian phylogenetic analyses were conducted using MrBayes version 3.2.6 (Huelsenbeck and Ronquist 2001) under the GTR+G nucleotide substitution model, selected using jModelTest (Posada 2008). Two independent runs were performed with four chains for 1,000,000 generations, with the first 25% discarded as burn-in. Posterior probabilities were used to assess node support.

To test for substitution saturation, Xia’s entropy-based index of substitution saturation (*I_ss_*) was computed for *cytb* sequences as well as separately for the first, second, and third codon positions (in DAMBE v7.5.53) (Xia et al. 2003; Pacheco et al. 2018b). The proportion of invariant sites was estimated for the sequences as a whole and for each codon position. *I_ss_* was compared to the critical value (*I_ss.c_*) at which phylogenetic information is insufficient (Table S2). Analysis was performed on fully resolved sites in the alignment only.

## Results

Blood samples were collected from 43 dogs, 30 cats and 106 wild, endemic carnivores from three species: *C. ferox* (n=27), *G. elegans* (n= 37), and *G. fasciata* (n=42) (Table S3). At the time of sampling, all animals appeared clinically healthy and did not show any pathological signs associated with infection by hemoparasites.

No ticks were detected on domestic dogs and cats while all 106 endemic carnivores were infested by ticks from *Haemaphysalis obtusa* (Ixodidae).

PCR tests from domestic and wild carnivores revealed that none of the dogs or cats were positive for piroplasm infection at the time of sampling whereas 91.5% (95%CI: 84.1-95.8%) wild carnivores were infected by piroplasmid. Table 2 shows the proportion of carnivores infected. All sequences obtained from parasites infecting the same carnivore species were identical. Unique sequences were deposited in GenBank under accession numbers QB246345-QB246348 for *18S rRNA* and QB250296-QB250299 for *cytb* genes. The parasite infecting *Galidictis fasciata* exhibited 90.2% and 92.5% *18S rRNA* sequence similarity with *Babesia* sp. sequences previously reported from wild lemurs (KT722785) and fruit bats (MH790142) in Madagascar, respectively (Larsen et al. 2016; Ranaivoson et al. 2019). The *18S rRNA* sequence from this parasite also showed 87% similarity with the sequences obtained from parasites infecting *G. elegans* and *Cryptoprocta ferox*.

**Table 2:** Prevalence of infection by piroplasmids in domestic and wild carnivorans living in and around protected areas of Madagascar. Numbers in square brackets indicate the 95% confidence interval of the prevalence estimates.

|  | n | Piroplasmid prevalence [95% Confidence interval] |
|---|---|---|
| Broad striped vontsira (*Galidictis fasciata*) | 42 | 93% [81% - 99%] |
| Ring-tailed vontsira (*Galidia elegans*) | 37 | 95% [82%-99%] |
| Fosa (*Cryptoprocta ferox*) | 27 | 85 % [65%-95%] |
| Betampona Natural Reserve (BET) | 18 | 83% [59%-96%] |
| Kirindy Mitea National Park (KMNP) | 9 | 89% [52-100%] |
| Domestic dogs (*Canis familiaris*) | 43 | 0% [0% - 10%] |
| Domestic cats (*Felis catus*) | 30 | 0% [0% - 14%] |

The parasites infecting *G. elegans* and *C. ferox* were identified as *Cytauxzoon* sp. Their 18S rRNA sequences showed 96% similarity to a sequence previously reported from meerkats (*Suricata suricatta*) in South Africa (KM025200; (Leclaire et al. 2015)). The sequence obtained from *C. ferox* in Betampona (BET) was 100% identical to that obtained from *C. ferox* in Kirindy Mitea National Park (KMN). Sequence similarity between the parasites infecting *G. elegans* and *C. ferox* was 98%.

The Bayesian phylogenetic analysis of the *18S rRNA* sequences (Figure 2) showed piroplasmid sequences obtained from Malagasy carnivores belonged to two distinct lineages. Sequences from *G. elegans* and *Cryptoprocta ferox* clustered within the *Cytauxzoon* Clade VII, forming a group distinct from *Cytauxzoon* sequences obtained from other mammalian hosts. In contrast, the sequences obtained from *G. fasciata* clustered within the *Babesia microti*-like Clade I, together with *Babesia* sequences from diverse mammalian hosts.

**Figure 2:**
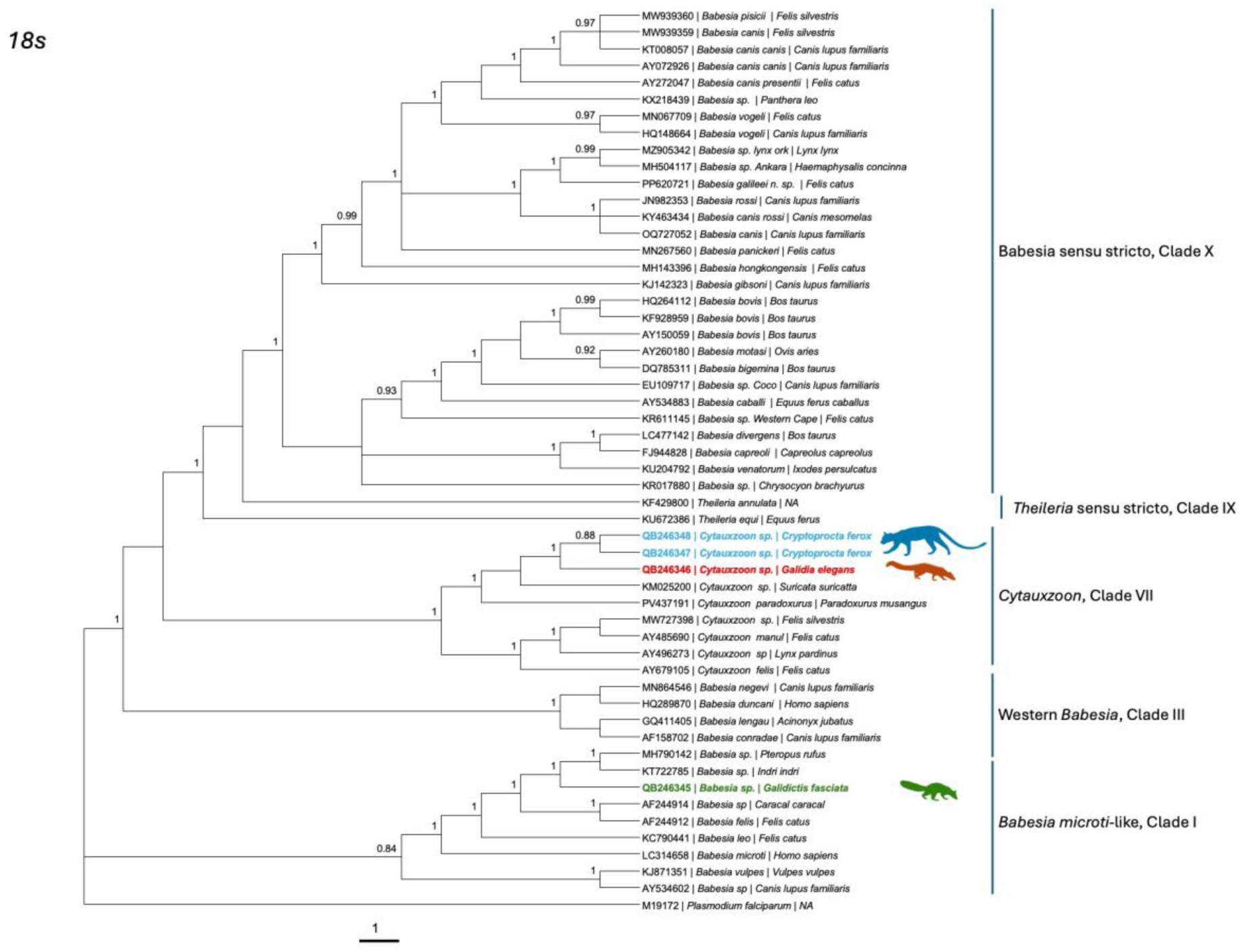
Bayesian phylogenetic tree of piroplasmid 18S rRNA sequences. The tree was inferred using the GTR+G substitution model in MrBayes from a 1,479-bp alignment of *18S rRNA* sequences. Posterior probability values are shown at nodes. Sequences generated in this study are highlighted by colored fonts. Piroplasmid sequences obtained from *Galidictis fasciata* (Green) cluster within the *Babesia microti*-like lineage (Clade I), whereas sequences obtained from *Galidia elegans* (Red) and *Cryptoprocta ferox* (Blue) cluster within the *Cytauxzoon* lineage (Clade VII). Representative sequences from GenBank are included for comparison, with host species indicated after sequence accession numbers. *Plasmodium falciparum* sequence was included as outgroup. Clade denomination following (Jalovecka et al. 2019).

The Bayesian phylogenetic analysis of the *cytb* gene sequences (Figure 3) of piroplasmids in Euplerids placed the parasites within a distinct clade comprising sequences from *G. elegans*, *G. fasciata*, and *C. ferox*. The *cytb* topology therefore supported genetic differentiation among piroplasmids infecting the three Malagasy euplerid hosts.

**Figure 3:**
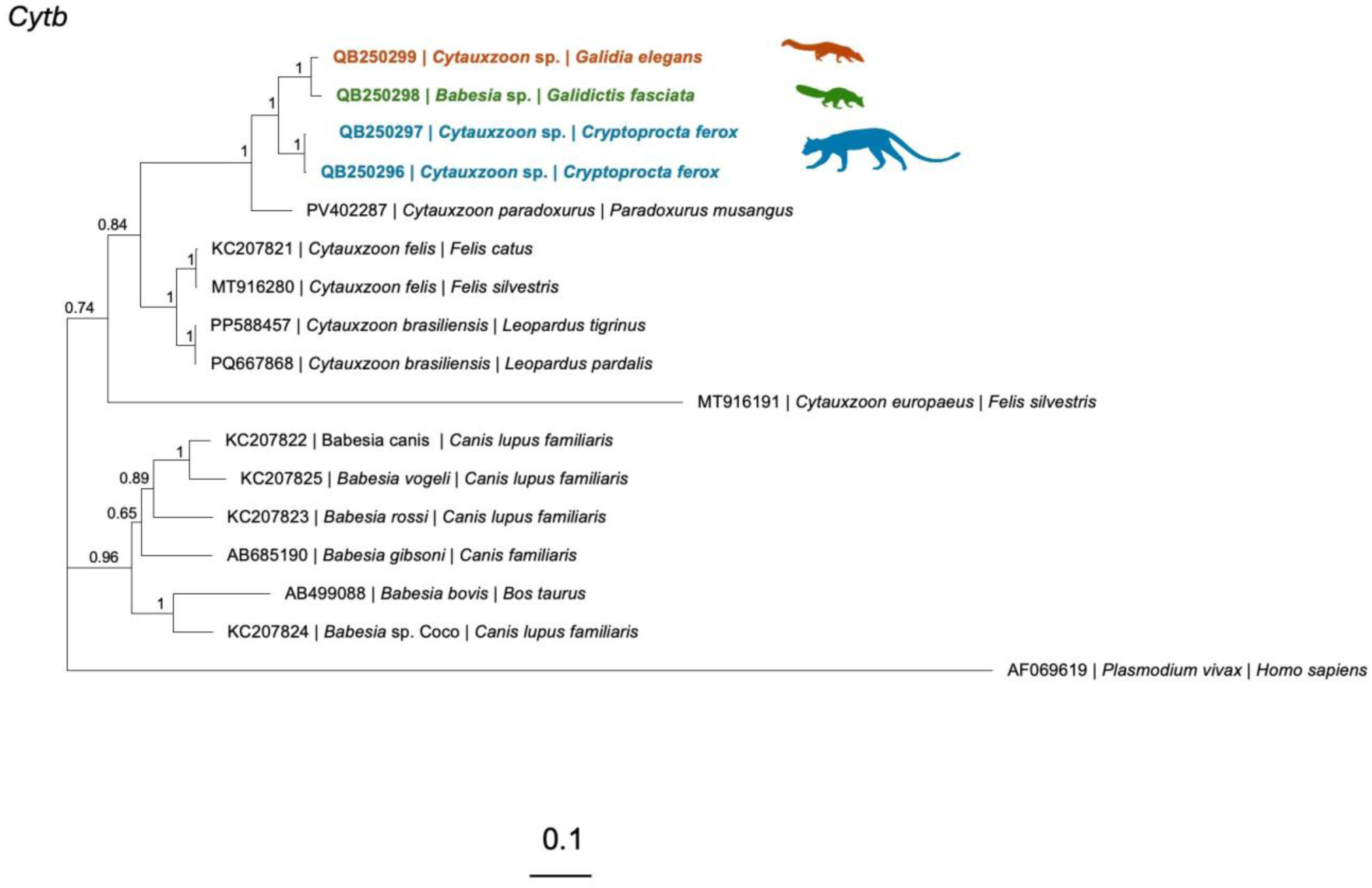
Bayesian phylogenetic tree of piroplasmid mitochondrial cytochrome b gene (*cytb*) sequences from Malagasy endemic carnivores and reference taxa. The phylogeny was inferred using Bayesian inference under the General Time Reversible (GTR) nucleotide substitution model. Sequences generated in this study are highlighted by host species: *Cytauxzoon* sp. from *Galidia elegans* (red), *Babesia* sp. from *Galidictis fasciata* (yellow), and *Cytauxzoon* sp. From *Cryptoprocta ferox* (green). Reference sequences are identified by GenBank accession number, parasite taxon, and host species. Posterior probability values are shown at nodes. Malagasy piroplasmid sequences form distinct host-associated lineages, with the *G. fasciata Babesia* sp. lineage occurring near the *G. elegans Cytauxzoon* sp. lineage. Branch lengths represent genetic divergence. *Plasmodium vivax* was included as an outgroup.

Testing for saturation of substitutions in the *cytb* sequences available for piroplasmids revealed little evidence of saturation, showing that the gene is phylogenetically informative. Xia’s indicator (*I_ss_*) was significantly below the threshold for saturation (*I_ss.c_*) when all, first, or second codon positions were considered (*p* < 0.01). We only observed evidence of saturation for third codon positions when an extremely asymmetrical tree was assumed (Table S2), indicating that *cytb* is unlikely to be saturated when comparing amongst piroplasmids.

## Discussion

We report piroplasmid parasites in wild Eupleridae, the endemic carnivore family of Madagascar. Specifically, we detected a high proportion (91.5%) of euplerids infected with piroplasmids and, more specifically, *Babesia* sp. infection in *G. fasciata* and *Cytauxzoon* sp. infection in *G. elegans* and *C. ferox*, whereas we did not detect either parasite genus in sympatric introduced carnivores. These findings expand the known host range of piroplasmids in Madagascar and suggest that infections are widespread among endemic carnivorans. Piroplasmid infections are common among wild carnivorans worldwide, although prevalence varies considerably among host species, geographic regions, and parasite genera (Alvarado-Rybak et al. 2016; Millán et al. 2024).

The high prevalence in Eupleridae suggests these hosts may play an important role in maintaining piroplasmids in the landscape. However, prevalence alone cannot establish reservoir competence, and experimental or longitudinal studies incorporating vector infection and transmission would be needed to determine whether these carnivorans serve as sources of infection for ticks. Persistent infections and/or frequent reinfection could contribute to the high prevalence observed in this population. The capacity of piroplasmids to persist through multiple tick developmental stages via transstadial transmission, together with prolonged or persistent infection in vertebrate hosts, may facilitate maintenance of these parasites within wildlife-vector systems (Bonnet et al. 2007; Chauvin et al. 2009; Jalovecka et al. 2019; Schnittger et al. 2022).

The absence of piroplasmid infection in samples from domestic dogs and cats may suggest that, in this environment, domestic and wild carnivores have little shared exposure (tick exchanges). However, studies show that domestic dogs and euplerids interact frequently in Madagascar and that ticks collected from endemic carnivorans can also infest domestic dogs. *Haemaphysalis obtusa*, the species of ixodid tick collected on euplerids in this study was previously detected from other endemic carnivores as well as the introduced *Viverricula indica,* and domestic dogs in various parts of the island (Uilenberg 1964; Apanaskevich and Goodman 2020). Alternatively, the apparent absence of piroplasmids in domestic carnivores may indicate that euplerid piroplasmids are unable to establish long-term infections in domestic dogs and cats, with introduced host species quickly clearing the parasite or succumbing to infection. For instance, *Babesia rossi*, the primary causative agent of canine babesiosis, can cause severe, often fatal disease in domestic dogs, whereas African wild canids may harbor the parasite with little or no apparent clinical disease (Penzhorn 2011). This contrast highlights the importance of host-parasite coevolution and susceptibility in determining the pathogenic consequences of piroplasmid infections. Further research is needed to assess the pathogenic potential of these parasites when they infect domestic dogs or other incidental or accidental hosts.

The Bayesian phylogenetic analysis based on the *18S rRNA* gene revealed that piroplasmid sequences obtained from Malagasy carnivores belonged to two distinct phylogenetic lineages. The 18S rRNA phylogeny (Figure 2) placed the *Babesia* sp. detected in *G. fasciata* within the *B. microti*-like group (Clade I), where it was most closely related to *Babesia* parasites previously reported from wild lemurs in Madagascar (Larsen et al. 2016) and fruit bats (Ranaivoson et al. 2019), rather than to *Babesia* reported from domestic cats and dogs elsewhere. This pattern suggests that the parasite infecting *G. fasciata* may be part of a lineage circulating among Malagasy wildlife rather than representing a recently introduced parasite associated with domestic carnivores. In contrast, the sequences detected in *Cryptoprocta ferox* and *G. elegans* showed closer phylogenetic affinities to *Cytauxzoon* parasites reported from carnivores outside Madagascar. These sequences were most closely related to *Cytauxzoon* sp. reported from meerkats in South Africa (KM025200) and *Cytauxzoon paradoxurus* (PV437191) detected in civets in Singapore (Leclaire et al. 2015; Chong et al. 2025). However, caution is warranted when interpreting 18S rRNA gene sequences because gene copy number and the extent of concerted evolution may vary among taxa, as shown for parasites from the genus *Plasmodium* as well as *Theileria equi* and *Babesia caballi* (Corredor and Enea 1993; Bhoora et al. 2009).

The *cytb* phylogeny further suggests that piroplasmids detected in Malagasy endemic carnivores comprise genetically distinct lineages potentially associated with their respective host species. All piroplasmid sequences obtained from Eupleridae clustered closely together, indicating low genetic divergence among parasites infecting these hosts and supporting their assignment to a distinct lineage. This pattern is consistent with hypothesis 1, local evolution and ancient diversification on the island of Madagascar, rather than spillover infections from domestic or introduced carnivores. However, broader sampling of piroplasmids from domestic and introduced carnivores, along with additional genetic markers, will be necessary to establish the extent of host association and evaluate potential cross-species transmission.

Phylogenetic analyses based on *18S rRNA* and mitochondrial *cytb* sequences revealed partial incongruence between gene trees (Figs. 2 and 3). Although the *18S rRNA* phylogeny placed the parasites infecting *G. fasciata* and those infecting *G. elegans* and *C. ferox* in separate genera, the *cytb* phylogeny grouped them into a single, well-supported clade distinct from other piroplasmid lineages. This discordance highlights uncertainty in their evolutionary relationships, and this interpretation should therefore be considered preliminary.

Mitochondrial markers such as *cytb* gene can be susceptible to phylogenetic artifacts, including long-branch attraction (Griffiths 1997; Bergsten 2005; Cuadrado et al. 2024). Conversely, although *18S rRNA* is widely used for piroplasmid phylogenetics, its multicopy nature may introduce additional complexity through intragenomic sequence variation and the potential for selective amplification of particular gene copies during PCR. Furthermore, differences in taxon representation between the two datasets complicate direct comparisons. The comparatively sparse representation of piroplasmid *cytb* sequences in public databases, relative to *18S rRNA*, further limits our ability to rule out sampling artifacts as a contributor to this pattern.

Resolving whether the unexpected clustering reflects shared ancestry or methodological artifacts will require multilocus analyses incorporating additional nuclear and mitochondrial markers, tests of sequence saturation and phylogenetic congruence, and expanded taxon sampling. In particular, sampling piroplasmids from both endemic and introduced Malagasy carnivores, as well as their associated tick vectors, would help clarify the evolutionary relationships and host associations of these parasites. Such efforts could also help determine whether the apparent absence of major piroplasmid lineages commonly associated with domestic dogs and cats in our dataset reflects a genuine lack of spillover into endemic carnivores or simply limited detection (Florin-Christensen et al. 2021; Singh et al. 2023).

## Acknowledgements

FTR, BLR were supported by the High Meadows Environmental Institute (HMEI), Princeton University. BLR was supported by the Princeton Catalysis Institute (PCI). FTR was supported by the Saint Louis Zoo WildCare Institute and Maxwell Hanrahan Foundation. The authors would like to acknowledge Dr. Hans Klompen and Dr. Dmitry Apanaskevitch for their input and thank Drs. Andriamihajarivo, Hertz Andrianalizah, Rebecca Lewis and Natacha Rasolozaka as well as the Ankoatsifaka Research Station, Madagascar Fauna and Flora Group and Madagascar National Parks for their support during the data collection.

## References

Alvarado-Rybak, M., L. Solano-Gallego, and J. Millán. 2016. A review of piroplasmid infections in wild carnivores worldwide: importance for domestic animal health and wildlife conservation. Parasites & Vectors 9:538.

Apanaskevich, D. A., and S. M. Goodman. 2020. Description of a new species of Haemaphysalis Koch, 1844 (Acari: Ixodidae) from the H. (Rhipistoma) asiatica subgroup, parasite of an endemic Malagasy carnivoran (Carnivora: Eupleridae). Systematic Parasitology:1–9.

Ardalan, A., M. C. Oskarsson, B. van Asch, E. Rabakonandriania, and P. Savolainen. 2015. African origin for Madagascan dogs revealed by mtDNA analysis. Royal Society Open Science 2:140552.

Bergsten, J. 2005. A review of long-branch attraction. Cladistics 21:163–193.

Bhoora, R. et al. 2009. Sequence heterogeneity in the 18S rRNA gene within Theileria equi and Babesia caballi from horses in South Africa. Veterinary Parasitology 159:112–120.

Bonnet, S. et al. 2007. Transstadial and transovarial persistence of Babesia divergens DNA in Ixodes ricinus ticks fed on infected blood in a new skin-feeding technique. Parasitology 134:197–207.

Chauvin, A., E. Moreau, S. Bonnet, O. Plantard, and L. Malandrin. 2009. Babesia and its hosts: adaptation to long-lasting interactions as a way to achieve efficient transmission. Veterinary Research 40:1–18.

Chong, S. Q. Y. et al. 2025. Cytauxzoon paradoxurus n. sp., a novel Cytauxzoon species identified in common palm civets in Singapore. Parasites & Vectors 18:175.

Corredor, V., and V. Enea. 1993. Plasmodial ribosomal RNA as phylogenetic probe: a cautionary note. Molecular biology and evolution 10:924–926.

Cuadrado, D. et al. 2024. Base-substitution rates of nuclear and mitochondrial genes for polyclad flatworms. Zoosystematics and Evolution 100:863–876.

Escalante, A. A., D. E. Freeland, W. E. Collins, and A. A. Lal. 1998. The evolution of primate malaria parasites based on the gene encoding cytochrome b from the linear mitochondrial genome. Proceedings of the National Academy of Sciences 95:8124–8129.

Farris, Z. J. et al. 2017. The times they are a changin’: Multi-year surveys reveal exotics replace native carnivores at a Madagascar rainforest site. Biological Conservation 206.

Florin-Christensen, M. et al. 2021. Pursuing effective vaccines against cattle diseases caused by apicomplexan protozoa. CABI Reviews 2021.

Florin-Christensen, M., and L. Schnittger. 2009. Piroplasmids and ticks: a long-lasting intimate relationship. Frontiers in Bioscience Volume:3064.

Griffiths, C. S. 1997. Correlation of Functional Domains and Rates of Nucleotide Substitution in Cytochromeb. Molecular Phylogenetics and Evolution 7:352–365.

Hixon, S. W. et al. 2021. Ecological Consequences of a Millennium of Introduced Dogs on Madagascar. Frontiers in Ecology and Evolution 9:689559.

Huelsenbeck, J. P., and F. Ronquist. 2001. MRBAYES: Bayesian inference of phylogenetic trees. Bioinformatics 17:754–755.

Jalovecka, M., D. Sojka, M. Ascencio, and L. Schnittger. 2019. Babesia Life Cycle – When Phylogeny Meets Biology. Trends in Parasitology 35:356–368.

Kjemtrup, A. M., and P. A. Conrad. 2000. Human babesiosis: an emerging tick-borne disease. International Journal for Parasitology 30:1323–1337.

Kjemtrup, A. M., J. Thomford, T. Robinson, and P. A. Conrad. 2000. Phylogenetic relationships of human and wildlife piroplasm isolates in the western United States inferred from the 18S nuclear small subunit RNA gene. Parasitology 120:487–493.

Lack, J. B., M. V. Reichard, and R. A. Bussche. 2012. Phylogeny and evolution of the Piroplasmida as inferred from 18S rRNA sequences. International Journal for Parasitology 42:353–363.

Larsen, P. A. et al. 2016. Blood transcriptomes reveal novel parasitic zoonoses circulating in Madagascar’s lemurs. Biology Letters 12:20150829.

Leclaire, S., S. Menard, and A. Berry. 2015. Molecular characterization of Babesia and Cytauxzoon species in wild South-African meerkats. Parasitology 142:543–548.

Leemans, I., D. Brown, P. Hooshmand-Rad, E. Kirvar, and A. Uggla. 1999. Infectivity and cross-immunity studies of Theileria lestoquardi and Theileria annulata in sheep and cattle: I. In vivo responses. Veterinary Parasitology 82:179–192.

Matjila, P., A. L. Leisewitz, F. Jongejan, H. J. Bertschinger, and B. L. Penzhorn. 2008. Molecular detection of Babesia rossi and Hepatozoon sp. in African wild dogs (Lycaon pictus) in South Africa. Veterinary Parasitology 157:123–127.

Millán, J. et al. 2024. Large-scale molecular survey for piroplasmids in Iberian wild carnivores. Parasitology Research 123:406.

Munson, L. et al. 2008. Climate Extremes Promote Fatal Co-Infections during Canine Distemper Epidemics in African Lions. PLoS ONE 3:e2545.

Nguyen, J. M. T., and S. Y. W. Ho. 2016. Mitochondrial rate variation among lineages of passerine birds. Journal of Avian Biology 47:690–696.

Oosthuizen, M. C., E. Zweygarth, N. E. Collins, M. Troskie, and B. L. Penzhorn. 2008. Identification of a Novel Babesia sp. from a Sable Antelope (Hippotragus niger Harris, 1838). Journal of Clinical Microbiology 46:2247–2251.

Pacheco, M. A. et al. 2018a. Primers targeting mitochondrial genes of avian haemosporidians: PCR detection and differential DNA amplification of parasites belonging to different genera. International Journal for Parasitology 48:657–670.

Pacheco, M. A. et al. 2018b. Mode and Rate of Evolution of Haemosporidian Mitochondrial Genomes: Timing the Radiation of Avian Parasites. Molecular Biology and Evolution 35:383–403.

Pacheco, M. A., and A. A. Escalante. 2023. Origin and diversity of malaria parasites and other Haemosporida. Trends in Parasitology 39:501–516.

Penzhorn, B. L. 2006. Babesiosis of wild carnivores and ungulates. Veterinary Parasitology 138:11–21.

Penzhorn, B. L. 2011. Why is Southern African canine babesiosis so virulent? An evolutionary perspective. Parasites & Vectors 4:51.

Posada, D. 2008. jModelTest: Phylogenetic Model Averaging. Molecular Biology and Evolution 25:1253–1256.

Ranaivoson, H. C., J.-M. Héraud, H. K. Goethert, S. R. Telford, L. Rabetafika, and C. E. Brook. 2019. Babesial infection in the Madagascan flying fox, Pteropus rufus É. Geoffroy, 1803. Parasites & Vectors 12:51.

Rasambainarivo, F., M. Andriamihajarivo, E. Dubovi, and P. G. Parker. 2018. Patterns of Exposure of Carnivores to Selected Pathogens in the Betampona Natural Reserve Landscape, Madagascar. Journal of wildlife diseases 54:386–391.

Rasambainarivo, F., Z. J. Farris, H. Andrianalizah, and P. G. Parker. 2017. Interactions Between Carnivores in Madagascar and the Risk of Disease Transmission. EcoHealth 14:691–703.

Rasambainarivo, F. T., J. Pomerantz, O. Rasolofoniaina, L. P. Rahajanirina, and L. Dollar. 2025. Physiological health benchmarks in free-ranging Fosa (Cryptoprocta ferox): establishing reference intervals to assess the impact of anthropogenic disturbance on an endemic apex predator. Journal of Mammalogy:gyaf063.

Sauther, M. L. et al. 2020. Taxonomic identification of Madagascar’s free-ranging “forest cats.” Conservation Genetics 21:443–451.

Schnittger, L., S. Ganzinelli, R. Bhoora, D. Omondi, A. M. Nijhof, and M. Florin-Christensen. 2022. The Piroplasmida Babesia, Cytauxzoon, and Theileria in farm and companion animals: species compilation, molecular phylogeny, and evolutionary insights. Parasitology Research 121:1207–1245.

Schnittger, L., A. E. Rodriguez, M. Florin-Christensen, and D. A. Morrison. 2012. Babesia: A world emerging. Infection, Genetics and Evolution 12:1788–1809.

Schreeg, M. E. et al. 2016. Mitochondrial Genome Sequences and Structures Aid in the Resolution of Piroplasmida phylogeny. PLoS ONE 11:e0165702.

Shock, B. C. et al. 2012. Variation in the ITS-1 and ITS-2 rRNA genomic regions of Cytauxzoon felis from bobcats and pumas in the eastern United States and comparison with sequences from domestic cats. Veterinary Parasitology 190:29–35.

Shock, B. C. et al. 2024. Description of B abesia coryicola sp. nov. from Florida pumas (Puma concolor coryi) from southern Florida, USA. International Journal for Parasitology: Parasites and Wildlife 24:100963.

Singh, P. et al. 2023. Babesia duncani multi-omics identifies virulence factors and drug targets. Nature Microbiology 8:845–859.

Soares, H. S. et al. 2017. Novel Anaplasma and Ehrlichia organisms infecting the wildlife of two regions of the Brazilian Amazon. Acta Tropica 174:82–87.

Solano-Gallego, L., Á. Sainz, X. Roura, A. Estrada-Peña, and G. Miró. 2016. A review of canine babesiosis: the European perspective. Parasites & Vectors 9:336.

Springer, A., C. Fichtel, S. Calvignac-Spencer, F. H. Leendertz, and P. M. Kappeler. 2015. Hemoparasites in a wild primate: Infection patterns suggest interaction of Plasmodium and Babesia in a lemur species. International Journal for Parasitology: Parasites and Wildlife 4:385–395.

Uilenberg, G. 1964. Notes sur les hématozoaires et tiques des animaux domestiques à Madagascar. Revue d’élevage et de médecine vétérinaire des pays tropicaux 17:337–359.

Veron, G. 2020. A review of the systematics of small mammalian carnivores. International Zoo Yearbook 54:35–42.

Veron, G., D. Dupré, A. P. Jennings, C. J. Gardner, A. Hassanin, and S. M. Goodman. 2017. New insights into the systematics of Malagasy mongoose-like carnivorans (Carnivora, Eupleridae, Galidiinae) based on mitochondrial and nuclear DNA sequences. Journal of Zoological Systematics and Evolutionary Research 55:250–264.

Wilson, E. B. 1927. Probable Inference, the Law of Succession, and Statistical Inference. Journal of the American Statistical Association 22:209–212.

Xia, X., Z. Xie, M. Salemi, L. Chen, and Y. Wang. 2003. An index of substitution saturation and its application. Molecular Phylogenetics and Evolution 26:1–7.

Yabsley, M. J., and B. C. Shock. 2012. Natural history of Zoonotic Babesia: Role of wildlife reservoirs. International Journal for Parasitology: Parasites and Wildlife 2:18–31.

Yoder, A. D. et al. 2003. Single origin of Malagasy Carnivora from an African ancestor. Nature 421:734–7.

Zintl, A., G. Mulcahy, H. E. Skerrett, S. M. Taylor, and J. S. Gray. 2003. Babesia divergens, a Bovine Blood Parasite of Veterinary and Zoonotic Importance. Clinical Microbiology Reviews 16:622–636.

